# Modeling HIV disease progression and transmission at population-level: The potential impact of modifying disease progression in HIV treatment programs

**DOI:** 10.1101/097337

**Authors:** Jennifer M. Ross, Roger Ying, Connie L. Celum, Jared M. Baeten, Katherine K. Thomas, Pamela M. Murnane, Heidi van Rooyen, James P Hughes, Ruanne Barnabas

## Abstract

**Introduction:** Mathematical models of HIV transmission that incorporate the dynamics of disease progression can estimate the potential impact of adjunctive strategies to antiretroviral therapy (ART) for HIV treatment and prevention. Suppressive treatment of HIV-positive persons co-infected with herpes simplex virus-2 (HSV-2) with valacyclovir, a medication directed against HSV-2, can lower HIV viral load, but the impact of valacyclovir on population HIV transmission has not been estimated.

**Methods:** We applied data on CD4 and viral load progression in ART-naïve persons studied in two HIV clinical trials to a novel, discrete-time Markov model. We validated our disease progression estimates using data from a trial of home-based HIV counseling and testing in KwaZulu-Natal, South Africa. Finally, we applied our disease progression estimates to a dynamic transmission model estimating the impact of providing valacyclovir to ART-naïve individuals to reduce onward transmission of HIV in three scenarios of different ART and valacyclovir population coverage. We assumed that valacyclovir reduced HIV viral load by 1.23 log copies/μL, and that persons treated with valacyclovir initiated ART more rapidly when their CD4 fell below 500 due to improved retention in pre-ART care.

**Results:** The average duration of HIV infection following acute infection was 9.5 years. The duration of disease after acute infection and before reaching CD4 200 cells/μL was 2.53 years longer for females than males. Relative to a baseline of community HIV testing and counseling and ART initiation at CD4 <=500 cells/μL, valacyclovir with increased linkage to care resulted in 166,000 fewer HIV infections over ten years, with an incremental cost-effectiveness ratio (ICER) of $4,696 per HIV infection averted. The Test and Treat scenario with 70% ART coverage and no valacyclovir resulted in 202,000 fewer HIV infections at an ICER of $6,579.

**Conclusion:** Even when compared with initiation of valacyclovir, a safe drug that reduces HIV viral load, universal treatment for HIV is the optimal strategy for averting new infections and increasing public health benefit. Universal HIV treatment should be pursued by all countries to most effectively and efficiently reduce the HIV burden.

## Introduction^1^

Dynamic HIV transmission models are used to guide implementation of HIV prevention and treatment interventions, estimate the cost and cost-effectiveness, and explore the potential impact of new strategies [1–3]. In principle, HIV models incorporate prevention interventions, such as behavioral changes, medications, or vaccines, based on known mechanisms of action or assumptions about how each intervention achieves its preventive effect. However, although HIV viral load is the primary predictor of HIV transmission, [4] few population modeling studies of HIV include a detailed description of the dynamics of HIV viral load along stages of HIV disease progression [5]. A potential modeling approach is nesting stochastic algorithms using Markov chain for HIV disease progression (i.e., changes in viral load and CD4 count over time) into HIV transmission models [6]. These estimates of changes in HIV viral load and CD4 count are particularly applicable when modeling interventions that exert a preventive effect by reducing HIV viral load.

While universal treatment with antiretroviral therapy (ART) is the most effective strategy for reducing HIV viral load and onward transmission[7, 8], 72% of HIV-positive people in western and central Africa and 46% of HIV-positive people in eastern and southern Africa lack access to ART [9], suggesting that adjunctive strategies to slow disease progression and prevent HIV transmission may be beneficial in these settings. Suppression of herpes simplex virus type two (HSV-2) in persons co-infected with HIV who are not yet on ART has been explored as an adjunctive tool for HIV prevention because HIV-HSV-2 co-infected individuals have faster CD4 count decline, increased HIV viral load and increased risk of onward transmission of HIV as compared to HIV mono-infected individuals [10, 11]. A previous study of the impact of HSV-2 suppression on HIV progression in co-infected individuals (the Partners in Prevention HSV/HIV Transmission Study) found that acyclovir 400mg, taken twice daily, did not reduce HIV transmission between HIV serodiscordant heterosexual couples, despite lowering viral load by 0.25 log copies/mL and reducing the occurrence of HSV-2 positive genital ulcers by 73% [11]. However, administration of twice daily high-dose (1.5g) valacyclovir, an acyclovir pro-drug, to HIV-HSV-2 co-infected individuals achieved a 76% HIV viral load reduction compared with treatment with acyclovir [12]. This finding raises the possibility that valacylovir could also impact HIV progression and transmission differently than acyclovir. Dynamic transmission modeling allows for estimation of how the additional reduction of HIV viral load achieved with valacyclovir could impact HIV progression, transmission, and at what programmatic cost.

Here, we report on a three step modeling investigation into the impact of co-infection treatment on HIV disease progression and transmission. First, we calibrate a discrete-time Markov model of HIV disease progression using data from the Partners in Prevention HSV/HIV Transmission Study [11] and the Partners PrEP Study [8] to assess CD4 and viral load changes over time in ART-naïve HIV-infected persons. Then, we validate our disease progression estimates using data on the distribution of CD4 counts and viral load from a trial of home-based HIV counseling and testing in KwaZulu-Natal, South Africa. Finally, we combine our disease progression model with a dynamic transmission model to compare strategies of valacyclovir HSV-2 suppression versus enhanced ART access to reduce onward transmission of HIV.

## Methods

### Study Population

Data to inform the model of HIV disease progression came from two studies of HIV prevention in serodiscordant heterosexual partnerships in sub-Saharan Africa—the Partners in Prevention HSV/HIV Transmission Study [11] and the Partners PrEP Study [8]. Briefly, the Partners HSV/HIV Transmission Study, a prospective placebo-controlled randomized study, enrolled 3,408 serodiscordant couples from eastern and southern Africa, in which the HIV-positive partner was ART-naïve and co-infected with HSV-2. The study evaluated the impact of HSV-2 suppression with acyclovir for the HIV-positive partner on HIV transmission [11]. The Partners PrEP Study, a prospective placebo-controlled randomized study, enrolled 4,758 serodiscordant couples from East Africa in which the HIV-infected partner was ART-naïve. While the primary objective of Partners PrEP was to evaluate the impact of pre-exposure prophylaxis for HIV-uninfected partners on HIV acquisition, CD4 and viral load were also measured at six-month intervals for individuals who acquired HIV infection. Nearly 56% of the HIV-negative partners were seropositive for HSV-2 at enrollment [8]. We included all CD4 and HIV viral load follow-up data for both studies from partners who were HIV-negative at enrollment and seroconverted during the study period.

### Data

CD4 count and viral load were measured at the beginning and end of all 6 and 12-month intervals post-seroconversion and used for this analysis. For intervals that ended with ART initiation, the CD4 count and VL were estimated by the aggregate distribution of CD4 and viral load measurements ≤3 months prior to ART initiation. There were 151 HIV seroconverters in the Partners in Prevention HSV/HIV Transmission Study and 138 HIV seroconverters in the Partners PrEP Study [8, 11].

### Analysis of Disease Progression

To estimate disease progression among HIV-infected individuals, we organized individuals into discrete CD4 and viral load categories. The amount of time spent in each CD4 and viral load category was then estimated by applying discrete-time Markov models for CD4 and viral load, and calculating the time to absorption (Fig. 1). Only observations with simultaneous CD4 and viral load measurements were included in the analysis. The proportions of individuals progressing from one CD4 and viral load category to another were assumed to form the transition matrix for progression from one category to another. For CD4 cell progression, we calculated the average time from CD4>500 cells/μL to absorption in each of the subsequent CD4 categories. The duration in each category was estimated to be the difference between times to absorption for adjacent absorption scenarios. A similar process was used for viral load progression, but with starting viral load of <1,000 copies/mL.

**Figure 1.**
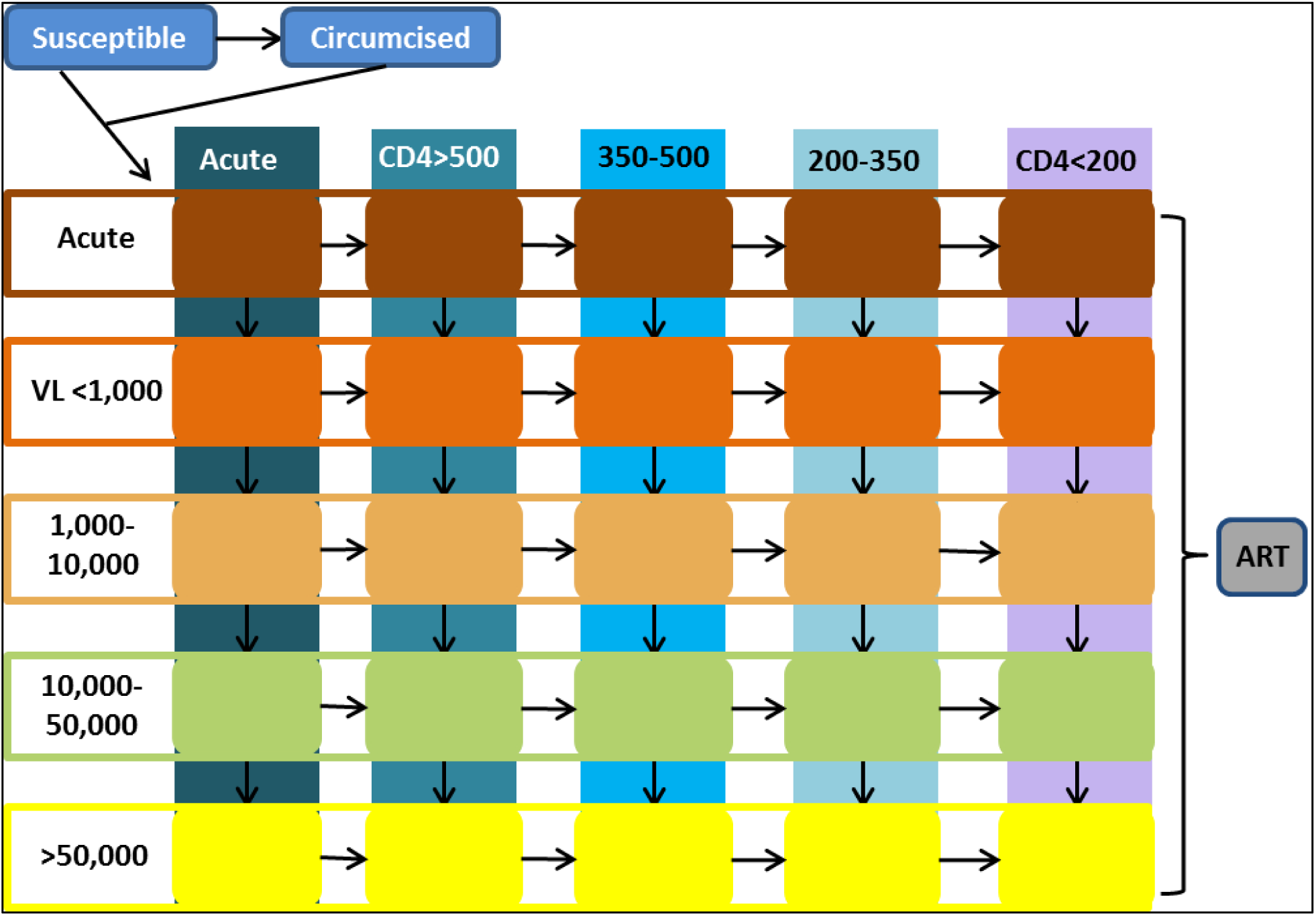
Model transition diagram. A diagram of the natural history of HIV infection. All movement is in one direction except for enrollment in and dropout from interventions from ART. Acute infection is modeled as a transient period immediately following HIV infection with high probability of onward HIV transmission, and is independent of CD4 count and viral load.

### Model Validation

We validated the disease progression model by comparing the annual cross-sectional distribution of CD4 counts and HIV viral load during the epidemic between our model and as observed values in KwaZulu-Natal from a previous study of home-based HIV testing and counseling (HTC) in KwaZulu-Natal, South Africa [13].

**Table 1.** Key parameters used in model. The parameters were based on the HBHCT study and other literature. For parameters with varying estimates, we used a value that best fit our model.

| Model Parameter | Value [Range] | Reference |
| --- | --- | --- |
| **Transmission Probability<sup>a</sup>** |  |  |
| Baseline Probability | 0.00053 | Boily *et al.*, Powers *et al.* [14, 15] |
| Acute | 26 x Baseline [0.0082, 0.015] | Hollingsworth *et al.* [16] |
| VL≤1,000 copies/mL | 1 x Baseline [0.01, 11] | Baeten *et al.* [17] |
| VL 1,000-10,000 copies/mL | 1.1 x Baseline [0.1, 3.8] | Baeten *et al.* [17] |
| VL 10,000-50,000 copies/mL | 3.1 x Baseline [1.3, 4.5] | Baeten *et al.* [17] |
| VL>50,000 copies/mL | 8.7 x Baseline [3.6-11.0] | Baeten *et al.* [17] |
| **ART Effectiveness for Reducing HIV Transmission<sup>b</sup>** | 90%<sup>c</sup> | Donnell *et al.*, Cohen *et al.* [7, 18] |
| **Baseline ART Coverage** | 36% of all HIV-infected persons | Barnabas *et al.* [19] |
| **HSV Prevalence** |  |  |
| Males | 60% | Barnabas *et al.* [20] |
| Females | 80% | Barnabas *et al.* [20] |
| **HSV Impact on HIV** |  |  |
| HIV acquisition in males | RR 2.8 | Barnabas *et al.* [20] |
| HIV acquisition in females | RR 3.4 | Barnabas *et al.* [20] |
| Disease progression | 20%<sup>d</sup> | Lingappa *et al.* [21] |
| **Costs** |  |  |
| HBHCT with Community Care Workers<sup>e</sup> | Per HIV-positive person: \$28.06<br>Per HIV-negative person: \$8.22 | Ying *et al.* [22] |
| ART | \$682 per person per year | CHAI [23] |
| Valacyclovir | \$571 per person per year | CHAI, personal communication |
| Hospitalization: pre-ART CD4≤200 cells/μL | \$121 per HIV-infected person per year | Meyer-Rath *et al.* [24] |
| Hospitalization: pre-ART CD4 200-350 cells/μL | \$58 per HIV-infected person per year | Meyer-Rath *et al.* [24] |
| Hospitalization: pre-ART CD4>350 cells/μL | \$39 per HIV-infected person per year | Meyer-Rath *et al.* [24] |
| Hospitalization: post-ART CD4 200-350 cells/μL | \$111 per HIV-infected person per year | Meyer-Rath *et al.* [24] |
| Hospitalization: post-ART CD4>350 cells/μL | \$45 per HIV-infected person per year | Meyer-Rath *et al.* [24] |

<sup>a</sup>Probability of HIV transmission per coital act assumes that HIV transmission is a Bernoulli process.
<sup>b</sup>Proportion of transmissions from or to a specific demographic averted due to intervention.
<sup>c</sup>Value includes 96% reduction in HIV transmission due to ART with 6% annual loss to follow-up.
<sup>d</sup>Individuals with HSV co-infection are 20% more likely to progress to the next CD4 stage.
<sup>e</sup>Community Care Workers (public-sector salary)

### HIV transmission model

We developed a dynamic, compartmental model of HIV transmission at population level in KwaZulu-Natal in which the disease progression model was embedded. The transmission mathematical model was parameterized from our observational home-based HTC study, which was conducted in Vulindlela, KwaZulu-Natal from September 2011 to May 2013 (Table 1). The model is stratified by age, gender, sexual risk, and HSV-2 status, with HIV-infected persons progressing through CD4 and viral load categories as defined for the Markov model. An individual’s HSV-2 status was HSV-2 uninfected, HSV-2 infected, or HSV-2 suppressed with valacyclovir prophylaxis. At baseline, we assume no valacyclovir prophylaxis is used and that 60% of HIV infected men are co-infected with HSV-2, and 80% of HIV-infected women are co-infected with HSV-2 [20]. HSV-2 infection is assumed to increase HIV transmission and HIV acquisition as estimated in a previous meta-analysis [20] (Table 1). The age-specific HIV incidence and prevalence were validated with independent South African national survey data [25].

### Valacyclovir intervention scenarios

We assessed the impact of valacyclovir prophylaxis on HIV progression and HIV transmission in three scenarios (Table 2). The baseline scenario assumes that valacyclovir is not provided, and that persons become eligible for ART when their CD4 falls below 500. The valacyclovir scenario assumes that 48% of individuals with CD4≤500 cells/μL and not using valacyclovir are on ART, with the initiation of valacyclovir leading to a 25% increase in ART coverage (60%), due to retention in pre-ART care and more rapid initiation of ART when an individual’s CD4 count falls below 500 cells/μL. The final scenario assumes a Test and Treat program where 70% of all HIV-positive individuals are on ART, and no one is taking valacyclovir prophylaxis. All scenarios assume that home-based HTC is a platform for reaching individuals, which reaches a greater proportion of individuals in a given community than strictly facility-based programs. We assumed that for HSV-2/HIV co-infected persons, valacyclovir prophylaxis reduces viral load by 1.23 log copies/mL, as observed previously, thus slowing CD4 progression [12]. In the first scenario, persons using valacyclovir were assumed to be on valacyclovir for five years, or until ART initiation.

**Table 2.** Intervention programmatic assumptions. The scenarios used in model to evaluate home-based HTC are based on an observational study of home-based HTC in KwaZulu-Natal from March 2011 to March 2013.

| Scenario | Drug Coverage by CD4 Category (cells/μL) | | | | Drug |
| --- | --- | --- | --- | --- | --- |
| | >500 after acute stage | 500 to 350 | 350 to 200 | ≤200 | |
| Baseline ART<sup>a</sup> | 0% | 20% | 60% | 90% | ART |
|  | 0% | 0% | 0% | 0% | Valacyclovir |
| Valacyclovir<sup>b</sup> | 0% | 60% | 60% | 90% | ART |
|  | 40% | 16% | 16% | 4% | Valacyclovir |
| Test and Treat<sup>c</sup> | 70% | 70% | 70% | 70% | ART |
|  | 0% | 0% | 0% | 0% | Valacyclovir |

<sup>a</sup>Baseline represents ART coverage with Home HTC.
<sup>b</sup>40% of HIV-positive persons not on ART use valacyclovir.
<sup>c</sup>Persons are eligible for ART at any disease stage, but only 70% of persons access ART.

Costs for this program were determined by estimates in the literature. The costs of ART and valacyclovir (1.5g) were estimated to be $682 [26] and $571 per person per year. HBHTC was assumed to cost $28.06 per HIV-positive person tested and $8.22 per HIV-negative person tested [22]. Finally, the costs of care for HIV-positive persons with and without ART were assumed to be as estimated by Meyer-Rath and colleagues[24].

## Results

### Sample Characteristics

The final sample included 5,388 6-month transitions and 10,706 12-month transitions, which were drawn from 289 unique individuals. Transitions are overlapping intervals of time that include movement of an individual from one CD4 and viral load compartment to another. The median length of follow-up following seroconversion was 15 months (range, 3 to 27 months) for the Partners HSV cohort and 31 months (range, 4 to 56 months) for the Partners PrEP cohort. The age distribution of seroconverters was similar between the two datasets, with 23% and 20% of individuals being less than 25 years of age in the Partners HSV and Partners PrEP cohorts, respectively, and 65% and 66% being less than 35 years, respectively.

### Estimated Duration of Disease Stage by CD4 and Viral Load

The overall durations of each disease stage by CD4 and viral load are shown in Table 3. Excluding acute infection, the average times spent with CD4>500 cells/μL, CD4 350–500 cells/μL, and CD4 200–350 cells/μL are 1.88 years, 1.22 years, and 5.90 years, respectively. The duration of disease after acute infection and before reaching CD4 200 cells/μL is 2.53 years longer for females than males. After assuming a three-month duration for acute infection and including estimates for mortality at each stage and for CD4≤200 cells/μL, overall life expectancy is estimated to be 11.58 years for females and 9.23 years for males.

Excluding acute infection, the average times spent with viral load ≤1,000 copies/mL, 1,000–10,000 copies/mL, and 10,000–50,000 copies/mL are 3.13 years, 1.99 years, and 4.40 years, respectively. The duration of disease after acute infection and before reaching viral load >50,000 copies/mL is 2.85 years longer for females than for males.

**Table 3.** Estimated duration of time in each CD4 (a) and Viral Load (b) category in ART-naïve persons, by gender. Estimates are based on average time to absorbing state in Markov model. Life expectancy estimate is the summation of time in each stage adjusted for CD4-specific mortality.

| (a) | Gender | Time (years) CD4 Stage |  |  |  |  | Life Expectancy<sup>a</sup> |
| --- | --- | --- | --- | --- | --- | --- | --- |
|  |  | Acute | >500 | 500 to 350 | 350 to 200 | ≤200 |  |
|  | Female | 0.25 | 1.94 | 1.35 | 6.71 | 1.96 | 11.58 |
|  | Male | 0.25 | 1.71 | 1.05 | 4.71 | 1.96 | 9.23 |
|  | Both | 0.25 | 1.88 | 1.22 | 5.90 | 1.96 | 10.66 |

| (b) | Gender | Time (years) Viral Load Stage |  |  |  |  | Life Expectancy<sup>a</sup> |
| --- | --- | --- | --- | --- | --- | --- | --- |
|  |  | Acute | ≤1,000 | 1,000 – 10,000 | 10,000 – 50,000 | >50,000 (est) |  |
|  | Female | 0.25 | 3.06 | 2.27 | 5.45 | 1.18 | 11.58 |
|  | Male | 0.25 | 3.44 | 1.45 | 3.04 | 1.50 | 9.23 |
|  | Both | 0.25 | 3.13 | 1.99 | 4.40 | 1.44 | 10.66 |

<sup>a</sup>Life expectancy estimates include mortality based on CD4.

### Validation of Disease Progression Estimates

With the disease progression estimates input as parameters in a mathematical model of heterosexual HIV transmission in KwaZulu-Natal, we determined the cross-sectional distribution of CD4 count and HIV viral load. We estimated that in 2012, the proportion of HIV-positive persons with CD4 counts <200, 200–350, 350–500, and >500 cells/μL were 12%, 38%, 19%, and 31%, which we found to be similar to estimates from KwaZulu-Natal, South Africa, of 11%, 22%, 25%, and 42%. The model also estimated the proportion of HIV-positive persons with viral load >50000, 10000–50000, 1000–10000, and <1000 copies/mL to be 22%, 23%, 12%, and 38%, which we found to be similar to estimates of 24%, 22%, 29%, and 25% (Fig. 2).

**Figure 2.**
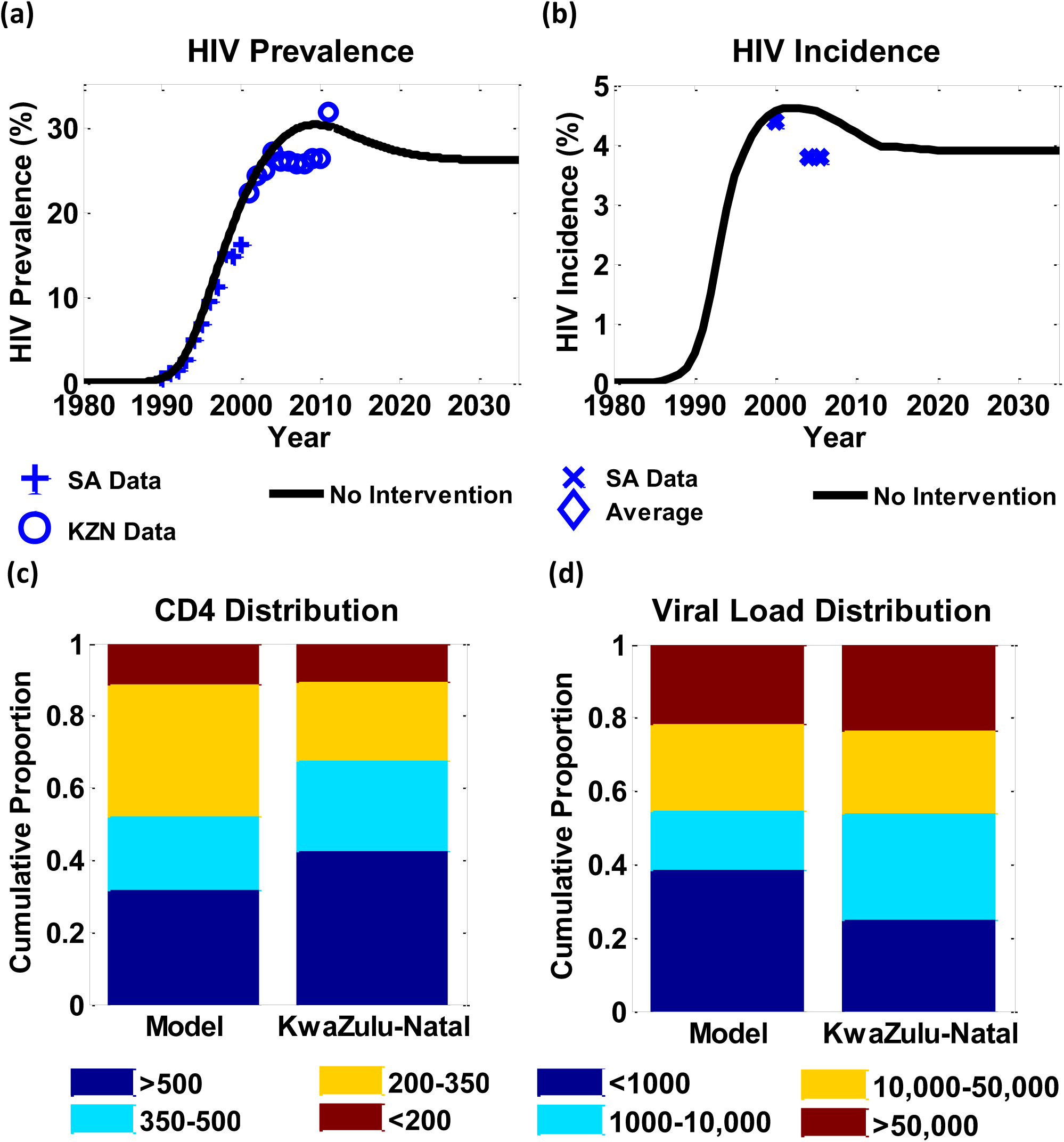
Model output for HIV prevalence and incidence. Model HIV prevalence (a) is similar to observed prevalence in KZN, and model HIV incidence (b) is similar to the average HIV incidence observed in KZN. Model output for the distribution of CD4 counts (c) and viral load (d) in HIV positive persons compared to the distribution seen in a study of Home HIV Testing and KwaZulu-Natal

### Estimated Impact and Cost-Effectiveness of Valacyclovir on HIV Disease Progression and Transmission

Relative to a baseline of community HTC, valacyclovir with increased linkage to care results in 166,000 fewer HIV infections over ten years, with an incremental cost-effectiveness ratio (ICER) of $4,696 per HIV infection averted (Table 4). The Test and Treat scenario of 70% ART coverage and no valacyclovir results in 202,000 fewer HIV infections at an ICER of $6,579, which is less than three times the per capita gross domestic product of South Africa and considered cost-effective.

Although valacyclovir is expected to prevent infections, it results in a reduction in quality-adjusted life-years (QALY) relative to baseline, due to slowing of CD4 decline and thus delaying ART eligibility in settings without universal eligibility for ART. The Test and Treat scenario, however, increases QALYs at an ICER of $570 per QALY gained relative to baseline.

**Table 4.**
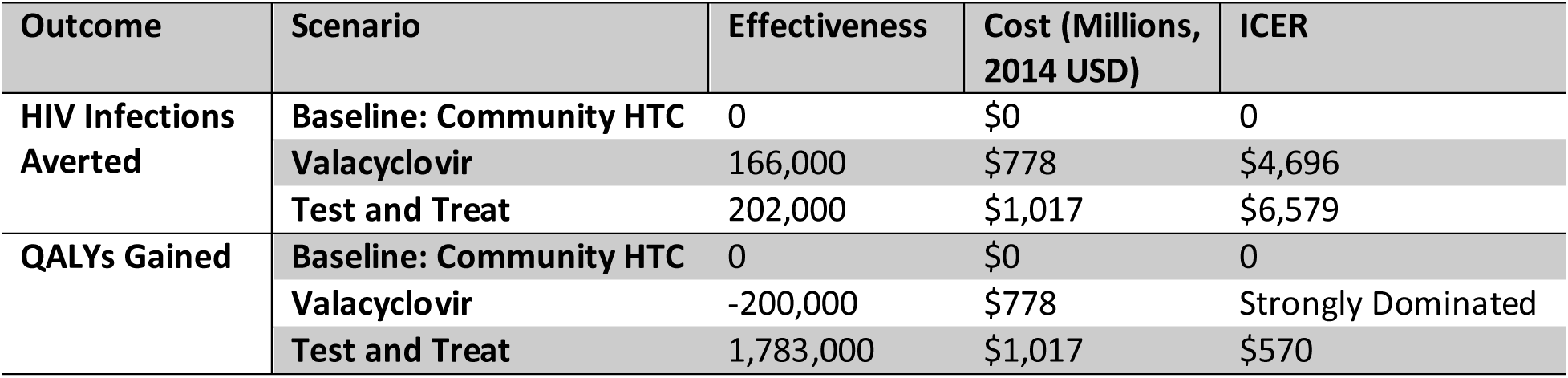
Effectiveness and cost-effectiveness of expanded ART coverage with and without valacyclovir prophylaxis. The baseline scenario assumes that 48% of HIV positive persons are virally suppressed. The outcomes reflect 3% annual discounting of health outcomes for a ten-year time horizon.

| Outcome | Scenario | Effectiveness | Cost (Millions, 2014 USD) | ICER |
| --- | --- | --- | --- | --- |
| HIV Infections Averted | Baseline: Community HTC | 0 | \$0 | 0 |
| | Valacyclovir | 166,000 | \$778 | \$4,696 |
| | Test and Treat | 202,000 | \$1,017 | \$6,579 |
| QALYs Gained | Baseline: Community HTC | 0 | \$0 | 0 |
| | Valacyclovir | -200,000 | \$778 | Strongly Dominated |
| | Test and Treat | 1,783,000 | \$1,017 | \$570 |

## Discussion

In this multi-part study, we first estimated HIV disease progression using data from two previous clinical trials fit to a discrete-time Markov model. Then, we validated and applied our disease progression findings to estimate the impact of valacyclovir prophylaxis taken by HIV-HSV-2 co-infected persons on HIV disease progression and transmission in a dynamic transmission model. In estimating disease progression, we found that women progress through the course of HIV disease more slowly than men, resulting in a longer life expectancy following seroconversion than men. Most of the additional time spent living with HIV for women is spent in the later stage of HIV infection (CD4 200 to 350 cells/μL and viral load 10,000 to 50,000 copies/mL). Estimating these HIV viral load values allowed us to further estimate the impact of providing valacyclovir to HIV-HSV-2 co-infected patients on onward transmission of HIV. Using our model of HIV transmission, we found that valacyclovir prophylaxis was a cost-effective strategy for averting HIV transmission and subsequent new infections. However, unintuitively the valacyclovir strategy reduced QALYs due to the slower CD4 decline and delay in ART eligibility among persons who were taking it. A test and treat scenario of 70% ART coverage without CD4 eligibility criteria was found to be the most cost-effective strategy.

### Disease progression modeling in context

Our modeled estimates of HIV disease progression are similar to those found in observational studies. In HIV-infected adults in Cote d’Ivoire and Uganda, the median time from seroconversion to CD4<350 cells/μL was 3.2 years, and from CD4<350 cells/μL to death was 7.6 years, which were similar to our estimates of 3.35 years and 7.86 years, respectively [27]. Two studies of European cohorts also found similar estimates. In the CASCADE Collaboration of primarily European individuals, estimates of 3.80 years from seroconversion to CD4<350 cells/μL and 7.10 years to CD4<200 cells/μL were similar to our estimates of 3.35 and 9.25 years, respectively [28]. The time from seroconversion to CD4<350 cells/μL was estimated to be shorter for Dutch MSM who acquired HIV infection in 2003–2007 versus in 1984–1995, with the estimate for 2003–2007 varying between 2.2 and 3.0 years between three different methods of calculation [29].

Previous observational studies have also found that women have slower disease progression than men. An analysis of the CASCADE Collaboration found that women have a relative risk of 0.76 of progressing to AIDS and a 0.68 relative risk of progressing to death relative to men [30]. Viral load has also been estimated to be lower in females than males at all CD4 levels in a cohort of intravenous drug users in the USA [31], as well as to increase at a slower rate in women than in men in an African cohort [32]. However, other studies have found no difference in CD4 count and progression between men and women [33]. Differences in results may be due to variable progression by HIV subtype [34, 35] or co-infection status [20].

### Valacyclovir prophylaxis in context

Despite prior published estimates of CD4 and viral load progression, few models have incorporated such estimates of disease progression into their assessments of the impact of valacyclovir therapy for HIV-HSV-2 co-infected persons. A previous study by Vickerman and colleagues estimated that acyclovir use in HIV-HSV-2 co-infected women, assuming increased retention in care for women prior to initiating ART, would cost $1130 per life-year gained [36], whereas the model used here found that valacyclovir prophylaxis would worsen health outcomes due to delayed ART initiation. The difference between our results can be attributed to different model structures. Whereas the model used by Vickerman and colleagues followed the life-time trajectory for 300 HIV-negative women, the model used here is population-level and measures outcomes on a ten-year time horizon. Furthermore, Vickerman et al. modeled the use of acyclovir rather than valacyclovir, with the former having been demonstrated to be less effective at reducing HIV viral load. Another model investigating the HIV-HSV-2 synergy was developed by Feng et al., who estimated the contribution of HSV-2 infection to the HIV epidemic, finding that nearly 10% of all HIV cases can be attributed to HSV-2 infection [37].

### Strengths and limitations of this study

A primary contribution of this study is developing a novel Markov model of HIV disease progression through CD4 and VL categories that generates cross-sectional CD4 and viral load prevalences which compare well with observed data. While the estimates were generally similar, the model did estimate that a larger proportion of the HIV-positive population would fall within the VL<1,000 category, relative to the observed data from KwaZulu-Natal (38% versus 25%), and a smaller proportion of the population to fall within the VL 1,000–10,000 category (12% versus 29%). This may be due to the structure of the model, where all individuals entered the VL <1000 category after acute infection. However, adjusting the modeled population viral load distribution to more closely match the KwaZulu-Natal data could be expected to further increase the benefit of the universal test and treat scenario, due to the efficacy of ART in reducing viral load and averting additional HIV transmissions. Additionally, although the data used to fit the Markov model were limited to recent HIV seroconverters rather than the general population of HIV positive persons, estimates of disease progression from seroconverters have been previously extrapolated to the general seroprevalent population [38].

Another strength of this analysis is that the datasets used to estimate the disease progression came from cohorts with high prevalence on HSV-2 seropositivity. These estimates would be generalizable to the many populations with high rates of HIV-HSV-2 co-infection [20], but may not apply as well to populations with low prevalence of HSV-2 infection. Finally, a major driver of our cost estimates were medication. Although our values were rigorously researched estimates, exact costs frequently change, particularly with costs decreasing in the near future.

### Future directions

As more national health programs adopt the WHO recommendation for universal ART eligibility, there will be additional opportunities to use models of disease progression to predict how expansion of ART in different care models will impact the epidemic. Understanding differences in disease progression by sex and by co-infection status will be important to include in these estimates. Models that explicitly include disease progression can be used to estimate the impact of therapeutic vaccines which aim to decrease the progression of HIV.

## Conclusion

Our finding that universal ART access achieves the greatest QALY gains supports current WHO goals of extending ART to all HIV-infected persons. Using validated estimates of disease progression in this model allowed us to estimate the population-level impact of a drug that slows disease progression. Even when compared with initiation of a safe drug (valacyclovir) that could potentially reduce HIV transmission in a setting of high HSV-2 prevalence, universal treatment for HIV is the optimal strategy for increasing QALYs and public health benefit. Universal HIV treatment should be pursued by all countries to most effectively and efficiently reduce the HIV burden.

## Acknowledgements

The authors acknowledge funding from the National Institutes of Health (NIH 5 R01 AI083034, 3 R0 AI083034-02S2, NIH Directors Award RC4 AI092552) and the University of Washington Center for AIDS Research (NIH P30 AI027757).

## Footnotes

1 Abbreviations – HSV-2, Herpes simplex virus type two; MSM, men who have sex with men; ART, antiretroviral therapy; VL, viral load; HTC, HIV testing and counseling; QALY, quality adjusted life year; ICER, incremental cost-effectiveness ratio; WHO, World Health Organization

